# Proteomic Analysis Uncovers Measles Virus Protein C Interaction with p65/iASPP/p53 Protein Complex

**DOI:** 10.1101/2020.05.08.084418

**Authors:** Alice Meignié, Chantal Combredet, Marc Santolini, István A. Kovács, Thibaut Douché, Quentin Giai Gianetto, Hyeju Eun, Mariette Matondo, Yves Jacob, Regis Grailhe, Frédéric Tangy, Anastassia V. Komarova

**Affiliations:** Viral Genomics and Vaccination Unit, Department of Virology, Institut Pasteur, CNRS UMR-3569, 75015 Paris, France; Université Paris Diderot, Sorbonne Paris Cité, Paris, France; Center for Research and Interdisciplinarity (CRI), Université de Paris, INSERM U1284; Network Science Institute and Department of Physics, Northeastern University, Boston, MA 02115, USA; Department of Physics and Astronomy, Northwestern University, Evanston, IL 60208-3109, USA; Department of Network and Data Science, Central European University, Budapest, H-1051, Hungary; Proteomics platform, Mass Spectrometry for Biology Unit (MSBio), Institut Pasteur, CNRS USR 2000, Paris, France; Bioinformatics and Biostatistics Hub, Computational Biology Department, Institut Pasteur, CNRS USR3756, Paris, France; Technology Development Platform, Institut Pasteur Korea, Seongnam-si, Republic of Korea; Laboratory of Molecular Genetics of RNA Viruses, Institut Pasteur, CNRS UMR-3569, 75015 Paris, France

**Author notes:** contributed equally to this work (Alice Meignié, Chantal Combredet).

## Abstract

Viruses manipulate central machineries of host cells to their advantage. They prevent host cell antiviral responses to create a favorable environment for their survival and propagation. Measles virus (MV) encodes two non-structural proteins MV-V and MV-C known to counteract the host interferon response and to regulate cell death pathways. Several molecular mechanisms underlining MV-V regulation of innate immunity and cell death pathways have been proposed, whereas MV-C host protein partners are less studied. We suggest that some cellular factors that are controlled by MV-C protein during viral replication could be components of innate immunity and the cell death pathways. To determine which host factors are targeted by MV-C, we captured both direct and indirect host protein partners of MV-C protein. For this, we used a strategy based on recombinant viruses expressing tagged viral proteins followed by affinity purification and a bottom-up mass spectrometry analysis. From the list of host proteins specifically interacting with MV-C protein in different cell lines we selected the host targets that belong to immunity and cell death pathways for further validation. Direct protein partners of MV-C were determined by applying protein complementation assay (PCA) and the bioluminescence resonance energy transfer (BRET) approach. As a result, we found that MV-C protein specifically interacts with p65/iASPP/p53 protein complex that controls both cell death and innate immunity pathways.

## INTRODUCTION

Measles virus (MV) is a member of the genus *Morbilliviru*s of the family *Paramyxoviridae*. This enveloped virus with a negative sense non-segmented RNA genome is responsible for measles, a childhood disease that used to cause 2.6 million deaths each year globally before the introduction in the 1970s of a live attenuated vaccine that reduced by 95% the incidence of the disease. Since, the MV vaccine has shown its safety and efficacy in over 2 billion children [1]. However, despite almost 50 years of vaccination history, we still know little about molecular mechanisms that make the attenuated MV vaccine so efficient. Furthermore, the measles vector platform is a promising plug-and-play vaccine platform technology for the rapid development of effective preventive vaccines against viral and other infectious diseases [1]. Another strong argument to study molecular mechanisms of MV replication and cellular pathways that are launched or controlled by MV is the efficacy of MV vaccine for oncolytic viral therapy [2]. Thus, numerous fundamental studies are needed to understand the principals of MV vaccine functioning as a preventive and therapeutic agent.

MV genome contains six genes encoding six structural proteins (N-P-M-F-H-L). The P gene encodes the P protein and two accessory proteins: V [3] and C [4]. Both MV-V and MV-C are viral virulence factors that exert multiple regulatory functions, suggesting numerous dynamic interactions with the human proteome. MV-V counteracts the host type-I IFN response by interacting with MDA5 (Melanoma-Differentiation-Associated protein 5), LGP2 (Laboratory of Genetics and Physiology 2), with Nuclear Factor-kappa B (NF-kB) subunits p65, IkB kinase α, STAT (Signal Transducer and Activator of Transcription) 1 and 2, JAK1 (Janus Kinase 1) and by interfering with their functions [5-8]. MV-V also interacts with p53 and p73 proteins, suggesting a link between MV-V and the cellular apoptosis pathway [9, 10]. By interacting with TRIpartite Motif containing protein 25 (TRIM25), MV-V stops the antiviral activities of RIG-I (retinoic-acid-inducible gene I) in the type-I IFN signaling pathway [11]. Numerous host protein partners have therefore been validated for MV-V, whereas MV-C network of interactions within MV-infected cells remains undetermined.

Viruses in the genera *Respirovirus, Henipavirus, and Morbillivirus* and *Tupaia paramyxovirus*-like viruses within the *Paramyxoviridae* family express one or more C proteins, which are relatively small basic proteins translated from the P mRNA in a different open frame from that for the P protein [12]. MV-C is a highly positively charged protein of 186 AA [4]. Using immunofluorescence, MV-C was localized in nucleus and in cytoplasmic inclusions [4, 13, 14]. MV-C possess a nuclear localization signal (NLS) and a nuclear export signal (NES), which allow MV-C to shuttle between the cytoplasm and the nucleus [14].

MV with abrogated expression of the C protein (MVΔC) has been evaluated *in vitro* and *in vivo* [15-23, 24]. In non-human primates MV-C can prevent cell death and is necessary for efficient viral replication as dramatically reduced expression of an MV antigen is detected in different tissues in comparison to *wt*MV [15, 24]. MV-C protein has been reported to inhibit the interferon (IFN-α/β/γ) responses [25-27]. MV-C interferes with the induction of IFN at the transcriptional level [28] and seems to control the induction of IFN by regulating viral RNA synthesis [21, 22, 29-31].

Numerous studies have demonstrated that viral defective interfering (DI) genomes are accumulated upon cell infection with MVΔC virus [21, 32]. These truncated replicative forms of the viral genome are directly linked to type-I IFN signaling through RIG-I (Retinoic acid-Inducible Gene-I) and protein kinase R (PKR) [22, 32-35]. Indeed, RIG-I is one of the viral RNA sensors triggering the production of pro-inflammatory cytokines, including IFN-α/β, and the establishment of an antiviral state in the host cells [36]. PKR is another sensor of viral dsRNA. Its activation triggers a cellular stress response leading to down-regulation of cellular protein synthesis due to the phosphorylation of eukaryotic translation initiation factor 2α (eIF2α) and to formation of stress granules [36]. In addition, the enhanced production of MV DI genomes by MVΔC that mediates PKR has correlated with a better activation of IRF-3 (IFN regulatory factor 3), NFκb and activating transcription factor 2 (ATF-2) that increase IFN production [33]. Thus, MV-C deletion from the virus suggests a direct contribution of MV-C on the viral RNA synthesis and either indirect or direct control of the host innate immune responses.

In cell culture, MV infection induces autophagy that is likely to favor the production of MV infectious particles and to delay the apoptosis induced by viral replication [23, 37]. MVΔC does not induce autophagy. Thus, it presents a defect in replication and is more apoptotic than the wild type virus [23, 38]. Former studies have also indicated that another paramyxovirus, Sendai virus, induces apoptosis, lacking expression of the C protein or containing a mutated C protein [39, 40]. These results suggest that MV replication induces apoptosis and that MV-C protein blocks this process *via* the modulation of virus replication process or *via* interaction with yet to be determined host factors.

To determine MV-C interaction network with host proteins, we captured both direct and indirect protein partners of MV-C protein expressed during the infection. For this, we used a strategy based on recombinant viruses expressing tagged viral protein followed by affinity purification and a bottom-up mass spectrometry-based proteomics analysis [10]. A list of specific MV-C protein-protein interactions (PPIs) was established in different cell lines, from which we validated MV-C direct PPI with proteins that belong to the immunity and cell death pathways. In particular, we uncovered MV-C direct PPIs with p65-iASPP-p53 protein complex that provide a mechanism on how MV-C protein can control both immunity and cell death responses.

## EXPERIMENTAL PROCEDURES

### Cells

HEK-293T (human embryonic kidney), HeLa (adenocarcinomic human epithelial cells), A549 (adenocarcinomic human alveolar basal epithelial cells) and Vero cells (African green monkey kidney cells) were maintained in DMEM, high glucose, GlutaMAX (#61965, Gibco, Switzerland) supplemented with 10% Fetal Bovine Serum (#10270, Gibco, Switzerland) and 10,000 units/mL of penicillin and 10,000 µg/mL of streptomycin (#15140122, Gibco, Switzerland) at 37°C in a 5% CO_2_ humidified atmosphere.

### Plasmid Constructions

for recombinant virus cloning construction of plasmids pTM-MVSchw, which contain an infectious MV cDNA corresponding to the anti-genome of the Schwartz MV vaccine strain has been described elsewhere [41].

To generate plasmid expressing a STrEP-tagged (WSHPQFEK) MV-C with an additional transcription unit (ATU) in between the H and L genes (ATU3, Fig. 1*A*), a two-step PCR-based strategy was used to produce coding sequences for MV-C and the Red Fluorescent Protein Cherry (CH) with either N- or C-terminal One-STrEP-tags. First, the One-STrEP-tag coding sequence was amplified by PCR from pEXPR-IBA105 using different primer pairs to add a flexible Tobacco Etch Virus protease linker (ENLYFQS) either at the N- or C-terminal to this sequence. In parallel, DNA fragments corresponding to MV-C or CH coding sequences were amplified using pTM3-MVSchw or pmCherry (Clontech, Palo Alto, USA) as a template. PCR products were finally combined and then amplified in a second PCR reaction to recover expected fused sequences, i.e. MV-C with either N-terminal or C-terminal One-STrEP tag, and CH with only N-terminal One-STrEP tag. These three amplicons contained unique *Bsi*WI and *Bss*HII sites at their extremities were first cloned in pCR2.1-TOPO plasmid (Invitrogen, USA) and Sanger sequenced. These plasmids were designated pTOPO/STrEP-C, pTOPO/C-STrEP, pTOPO/STrEP-CH. Finally, sequences were introduced in the ATU3 of pTM3-MVSchw vector [41] after *Bsi*WI/*Bss*HII digestion. The resulting plasmids were designated pTM3-MV/STrEP-C, pTM3-MV/C-STrEP, and pTM3-MV/STrEP-CH. All MV insertions respect the “rule of six,” which stipulates that the number of nucleotides of MV genome must be a multiple of six [42]. Recombinant viruses were rescued, and virus titers and single-step growth curves were determined as previously described [41].

**Fig. 1.**
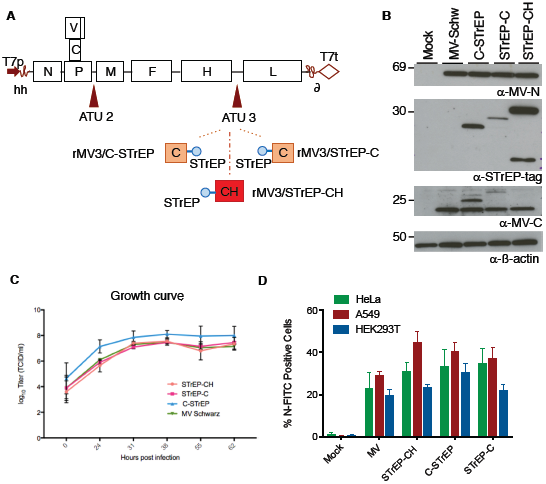
Recombinant viruses express tagged MV-C proteins, and replicate at high titers. *A*. Schematic representation of recombinant virus genomes. MV negative-sense RNA genome is displayed with its 3’ end on the left, with the six genes indicated by capital letters and depicted as rectangles. The additional transcriptional units encoding for One-STrEP-tagged MV-C (STrEP-C or C-STrEP) or CH protein (STrEP-CH) are inserted between H and L genes. The blue oval represents the One-STrEP-tag sequence. T7p is a T7 RNA polymerase promoter sequence, hh is a hammerhead ribozyme, T7t is a T7 RNA Polymerase termination signal, d is a hepatitis delta ribozyme. *B*. HeLa cells were infected with the native MVSchw strain, or the rMV2/STrEP-C and rMV2/C-STrEP expressing the One-STrEP-tagged MV-C protein or the rMV2/STrEP-CH expressing the One-STrEP-tagged mCherry (CH) protein. Expressions of native and One-STrEP-tagged MV-C proteins was determined by western blot using anti-C polyclonal and anti-STrEP tag monoclonal antibodies, respectively. Anti-MV-N antibody and anti-β−actin served as controls for Measles infection and loading, respectively. *C*. Virus growth curves obtained for rMV2/STrEP-C and rMV2/C-STrEP and rMV2/STrEP-CH. MV Schwarz strain was used as a control. Vero cells were infected with MV Schwarz, rMV2/STrEP-C and rMV2/C-STrEP, or rMV2/STrEP-CH at MOI of 0.1. Cell-associated virions were recovered at each time point, and titers were determined using the TCID50 method. Two biological experiments were performed and each point represent the mean and standard deviation of the two values. *D*. The efficiency of tagged recombinant viruses replication in HEK293T, HeLa, and A549 cells analyzed by FACS. Cells were infected at MOI1. 24h post-infection cells were harvested, fixed and stained using an anti-N antibody to measure percentage of N positive cells. Experiments were performed two times and data represent means ± SD of the technical triplicates of the most representative experiment.

For Protein Complementation Assay (PCA) based on the split luciferase (N2H) vectors pSNL-N1 and pSNL-N2 expressing the Nluc1 and Nluc2 complementary fragments of Nanoluciferase (Nluc) linked to the N-terminally of tested proteins were used [43]. The ORFs encoding for the selected proteins (Supp.Table 1) cloned into vector pDON223 (Gateway system; Invitrogen, USA) by recombination cloning (Gateway system; Invitrogen, USA) were collected from human ORFeome Collection v8.1 (generous gift from CCSB, DFCI, Harvard medical school, Boston). The resulting entry clones were then transferred into Gateway-compatible PCA destination vector pSNL-N1. The resulting pSNL-N1-”C interactors” were sequenced using forward primer (5’-GCTGAAGATCGACATCCATGTC-3’). The sequences were compared to the human genome by performing a blast search to identify the cDNA and the isoform when it is known. The in-frame fusion with the Nluc1 fragment of Nluc was verified. The ORFs encoding the MV-C protein and Muc7 protein (Mucin 7 was used as a negative control) were cloned into entry vector pDON207 (Gateway system; Invitrogen, USA). They were transferred into the destination vector pSNL-N2 by Gateway cloning. The resulting pSNL-N2-C and pSNL-N2-Muc7 were sequenced using forward primer (5’-CGGAGTGACCGGCTGGCGGCTG-3’).

For Bioluminescence Resonance Energy Transfer (BRET) assay we used the pEYFP-C1/N1 plasmids encompassing EYFP tag in the N- or C-terminal positions (Clontech, Mountain View, CA). To obtain pNluc-C1/N1 plasmids, the pEYFP-C1/N1 plasmids were modified by replacing the EYFP to the Nano-Luciferase (Nluc) coding sequence. A simple PCR amplification strategy was used to clone MV-C and p53 coding sequences in either pNluc-C1/N1 or pEYFP-N1 expression vectors. p53 was amplified from spleen cDNA library (Invitrogen, USA). For MV-C was amplified from pTM-Schwarz using different primers: forward 5’-TTCGAATTCTATGTCAAAAACGGA and reverse 5’-TAAGCGCGCGTCGACTCAGGAGCTCGTGGAT, digested with *Eco*RI and *Sal*I DNA restriction endonucleases and cloned in pNluc-C1 vector. A similar protocol was used to obtain pNluc-N1-MVC plasmid, except that primers were forward 5’-TTAGAATTCGCCACCATGTCAAAAACGGAC and reverse 5’-TAAGTCGACTGGGAGCTCGTCGTGGA and the vector was pNluc-N1. The same protocol was repeated to obtain pEYFP-N1-p53 plasmid encoding p53 with forward 5’-TTAGAATTCATGGAGCCGCAGTCAGA and reverse 5’-AATGTCGACGTCTGAGTCAGGCCCTTCTG primers. The resulting plasmids were designated as Nluc-C, C-Nluc, YFP-p53. DAXX-YFP and iASPP-YFP were obtained by recombination cloning (Gateway system). pDON223-DAXX and pDON223-iASPP were then transferred into Gateway-compatible destination vector pEYFP-C1. Plasmids were Sanger sequenced using CMV promoter primer. pYFP-p65 which contain EYFP reporter gene fused to p65 was a king gift of Dr.Hervé Bourhy from Lyssavirus Epidemiology and Neuropathology Laboratory of Institut Pasteur [44]. BRET negative control plasmids cSOD1-YFP, SOD1-Nluc and YFP-Nluc have been described [45].

### Viruses

The MV Schwarz vaccine strain (MVSchw, GenBank accession no. AF266291.1) was described in [41]. rTM-MVSchwΔC, rTM3-MV/STrEP-C, rTM3-MV/C-STrEP, and TM3-MV/STrEP-CH, rTM-MVSchwΔC-luc plasmids were used to rescue corresponding viruses using the helper-cell-based rescue system described in [41].

### Antibodies

Rabbit polyclonal anti-MV-C (kindly provided by Dr. Kaoru Takeuchi [46]), a rabbit polyclonal NF-kB p65 antibody (ab16502, Abcam, USA), a mouse monoclonal p53 Antibody (PAb 240, Invitrogen, USA), a mouse monoclonal antibody DAXX (sc-8043, SantaCruz, USA), a mouse monoclonal antibody iASPP (sc-398566, SantaCruz, USA), a mouse anti-p53 (556534,BD Biosciences)a mouse anti-N mAb (clone 25; kindly provided by Pr. Chantal Rabourdin-Combe, [47]), a monoclonal mouse Strep-tag antibodies (#34850, Qiagen), a monoclonal anti-actin antibody (A5441, Sigma-Aldrich, USA). As secondary antibodies, we used for western blot – a goat anti-mouse immunoglobulins HRP conjugated (P0447, Dako, Germany) or anti-rabbit (P0399, Dako, Germany); for immunofluorescence - a goat anti-mouse IgG (H+L) secondary antibody Alexa Fluor 555 (#A31572, Invitrogen, USA), a goat anti-rabbit IgG (H+L) highly cross-adsorbed secondary antibody Alexa Fluor 647 (#A21240, Invitrogen, USA); for flow cytometry analysis – an anti-Measles antibody, nucleoprotein, clone 83KKII, FITC-conjugated (#MAB8906F, Sigma-Aldrich, USA).

### Western Blots Assays

A549, HeLa or HEK293T cells were mock infected (treated with media alone) or infected with MV recombinant viruses at a multiplicity of infection (MOI) of 1. Protein lysates were fractionated by SDS-PAGE gel electrophoresis on 4–12% (or 12%) Nupage Bis–Tris gels with MOPS running buffer (#161-0788, Invitrogen, USA) and transferred to cellulose membranes (#10600016, GE Healthcare, USA). Membrane was incubated with antibodies. Peroxidase activity was visualized with SuperSignal West Pico PLUS Chemiluminescent Substrate (#34580, Thermo Fisher Scientific, USA).

### Flow Cytometry

HEK293T, HeLa, and A549 (2×105 cells per well) were plated in a 24-well plate and infected at MOI 1 with the corresponding recombinant viruses. Cells were washed twice with Dulbecco’s phosphate-buffered saline (PBS, #14190, Invitrogen, USA) and 2% foetal calf serum (FCS) and then fixed in PBS containing 4% paraformaldehyde. Cells were permeabilized with PermWash buffer (#554723, BD Biosciences, Germany), incubated with the primary antibody anti-N at 4°C for 30 min, washed in PermWash and incubated with the secondary antibody anti-mouse. Cells were washed twice with PBS 2% FCS before analysis by flow cytometry using a MACSQuant cytometer (Miltenyi Biotec, Germany) and analysis was done with the software FlowJo (vers 7.6).

### Affinity Purification of MV-C-specific Protein Complexes

HEK293T, HeLa, and A549 (2×107 cells per well) were mock infected (treated with media alone) or infected at an MOI of 1 for 24 h with MV-Schwarz, rTM3-MV/C-STrEP, rTM3-MV/STrEP-C and rTM3-MV/STrEP-CH. Cells were washed twice with cold PBS and lysed in 4 ml of lysis buffer (20mM MOPS-KOH pH7.4, 120mM of KCl, 0.5% Igepal, 2mM β-Mercaptoethanol), supplemented with Complete EDTA-free Protease Inhibitor Cocktail (#11836170001, Roche Diagnostics, USA). Cell lysates were incubated on ice for 20min with gentle mixing every 5 min, and then clarified by centrifugation at 16,000g for 15min at 4°C. The clarified cell lysates were incubated for 2 h on a spinning wheel at 4°C with 100µl of StrepTactin Sepharose High Performance beads (#28935599, GE Healthcare, USA). Beads were collected by centrifugation (1600 g for 5 min at 4°C) and washed twice for 5 min on a spinning wheel with 5ml of washing buffer (20mM MOPS-KOH pH7.4, 120mM of KCl, 2mM β-Mercaptoethanol) supplemented with Complete Protease Inhibitor Cocktail. Precipitates were eluted using 200µl Strep-Tactin Elution Buffer with Biotin (#2-1019-025, IBA, Germany). Finally, eluted proteins were precipitated overnight at 4°C with TCA (12% final concentration). Protein pellets were washed twice with ice-cold acetone, and resuspended in Urea 8M, 4%SDS final solution or were either analysed by western blotting or by LC-MS/MS.

### LC-MS/MS Analysis

For MV-C-specific protein co-complex (OUTPUTS), the digestion of the protein pellets were resuspended in 45µL of a Urea 8M / NH4HCO3 100mM denaturation buffer and sonicated 2 x 1min on ice. Cysteine bonds were reduced with 50mM TCEP (#646547, Sigma-Aldrich, USA) for 1h and alkylated with 50mM iodoacetamide (#I114, Sigma-Aldrich, USA) for 1h at room temperature in the dark. Samples were digested with rLys-C (#V1671, Promega, France) ratio 50:1 (protein:rLysC) for 3h/37°C and then digested with Sequencing Grade Modified Trypsin (#V5111, Promega, France) ratio 50:1 (protein:trypsin) overnight at 37°C. The digestion was stopped with 4% formic acid (FA) (#94318, Fluka) and digested peptides were purified with C18 Spin Columns Pierce™ (#89870, ThermoFisher Scientific, USA). Peptides were eluted with 2 x 80% Acetonitrile (ACN) / 0.1% FA. Resulting peptides were speedvac dried and resuspended in 2% Acetonitrile / 0.1% FA.

For input samples digestion, clarified cell lysates were processed as described in Erde et al. [48]. Briefly, 30 000 MWCO centrifugal units (Amicon^®^ Centrifugal Filters, Merck) and collection tubes were passivated in 5% (v/v) TWEEN^®^-20 and all subsequent centrifugation steps were carried out at 14 000g for 10 min. For each sample, 50µg of proteins were transferred into a filter unit and lysis buffer was exchanged with an exchanged buffer composed of 8 M urea, 0.2% DCA, 100 mM ammonium bicarbonate pH8.0. Disulfide bonds were reduced with 5mM TCEP for 1h and then alkylated with 50 mM iodoacetamide in the dark for 1h. Finally, one exchange buffer and two washes with the digestion buffer (0,2 % DCA / 50 mM ammonium bicarbonate pH 8) were performed before adding 100 μL of digestion buffer containing 1:50 ratio of sequencing-grade modified trypsin. Proteolysis was carried out at 37 °C overnight and peptide recovery was performed with 50 mM ammonium bicarbonate pH8.0. DCA was removed by acidification and phase transfer. Finally, dried peptides were resuspended in 2% acetonitrile, 0.1% FA prior to LC-MS/MS analysis.

LC-MS/MS analysis of MV-C specific protein complexes was performed on a Q ExactiveTM Plus Mass Spectrometer (Thermo Fisher Scientific, USA) coupled with a Proxeon EASY-nLC 1000 (Thermo Fisher Scientific, USA). 350ng of peptides were injected onto a home-made 50 cm C18 column (1.9 μm particles, 100 Å pore size, ReproSil-Pur Basic C18, Dr. Maisch GmbH, Ammerbuch-Entringen, Germany) and eluted with a multi-step gradient from 2 to 27% ACN in 105min, 27 to 50% ACN in 40min and 50 to 60% ACN in 10min, at a flow rate of 250 nL/min over 185 min. Column temperature was set to 60°C. MS data were acquired using Xcalibur software using a data-dependent method. MS scans were acquired at a resolution of 70,000 and MS/MS scans (fixed first mass 100 m/z) at a resolution of 17,500. The AGC target and maximum injection time for the survey scans and the MS/MS scans were set to 3E6, 20ms and 1E6, 60ms respectively. An automatic selection of the 10 most intense precursor ions was activated (Top 10) with a 45s dynamic exclusion. The isolation window was set to 1.6 m/z and normalized collision energy fixed to 28 for HCD fragmentation. We used an underfill ratio of 1.0% for an intensity threshold of 1.7E5. Unassigned precursor ion charge states as well as 1, 7, 8 and >8 charged states were rejected and peptide match was disabled.

LC-MS/MS analysis of clarified total lysates (INPUTS) was performed on a Q ExactiveTM Plus Mass Spectrometer (Thermo Fisher Scientific, USA) coupled with a Proxeon EASY-nLC 1200 (Thermo Fisher Scientific, USA). 1µg of peptides were injected onto a home-made 50 cm C18 column (1.9 μm particles, 100 Å pore size, ReproSil-Pur Basic C18, Dr. Maisch GmbH, Ammerbuch-Entringen, Germany) and eluted with a multi-step gradient from 3 to 29% buffer B (80% ACN) in 135min, 29 to 56% buffer B in 20min, at a flow rate of 250 nL/min over 182 min. Column temperature was set to 60°C. MS data were acquired using Xcalibur software using a data-dependent method. MS scans were acquired at a resolution of 70,000 and MS/MS scans (fixed first mass 100 m/z) at a resolution of 17,500. The AGC target and maximum injection time for the survey scans and the MS/MS scans were set to 3E6, 20ms and 1E6, 60ms respectively. An automatic selection of the 10 most intense precursor ions was activated (Top 10) with a 45 s dynamic exclusion. The isolation window was set to 1.6 m/z and normalized collision energy fixed to 28 for HCD fragmentation. We used a minimum AGC target of 1.0E4 corresponding to an intensity threshold of 1.7E5. Unassigned precursor ion charge states as well as 1, 7, 8 and >8 charged states were rejected and peptide match was disabled.

### Bioinformatics Analysis of LC-MS/MS Data

Raw data were analysed using MaxQuant software version 1.5.0.30 [49] for output samples and version 1.5.1.2 for inputs using the Andromeda search engine [50]. The MS/MS spectra were searched against two databases: the Human SwissProt database (20,203 entries from UniProt the 18/08/2015) and the Morbillivirus SwissProt database (90 entries from UniProt the 12/01/2016). Variable modifications (methionine oxidation and N-terminal acetylation) and fixed modification (cysteine carbamidomethylation) were set for the search and trypsin with a maximum of two missed cleavages were chosen for searching. The minimum peptide length was set to 7 amino acids and the false discovery rate (FDR) for peptide and protein identification was set to 0.01. At least a unique peptide per protein group was required for the identification of the proteins. The main search peptide tolerance was set to 4.5 ppm and to 20 ppm for the MS/MS match tolerance. Second peptides were enabled to identify co-fragmentation events and match between runs option selected with a match time window of 0.7 min for an alignment time window of 20 min. Quantification was performed using the XIC-based LFQ algorithm with the Fast LFQ mode as previously described [51]. Unique and razor peptides, included modified peptides, with at least 2 ratio counts were accepted for quantification. Respectively to the three different studied cell lines (HeLa, A549 and HEK 293T cell lines), three MaxQuant analysis were computed and analysed separately.

### Statistical Analysis

For the differential analyses, proteins identified in the reverse and contaminant databases and proteins “only identified by site” were first discarded from the list of identified proteins. Then, only proteins with three quantified intensity values in a condition were kept. After log2 transformation of the leftover proteins, LFQ values were normalized by median centering within conditions (normalizeD function of the R package DAPAR [52]). Remaining proteins without any LFQ value in one of both conditions have been considered as proteins quantitatively present in a condition and absent in another. They have therefore been set aside and considered as differentially abundant proteins. Next, missing values were imputed using the imp.norm function of the R package norm (Novo A. A. norm: Analysis of multivariate normal datasets with missing values. 2013 R package version 1.0-9.5). Proteins with a fold-change under 2.0 have been considered not significantly differentially abundant. Statistical testing of the remaining proteins (having a fold-change over 2.0) was conducted using a limma t-test [53] thanks to the R package limma [54]. An adaptive Benjamini-Hochberg procedure was applied on the resulting p-values thanks to the function adjust.p of R package cp4p [55] using the robust method of [54] to estimate the proportion of true null hypotheses among the set of statistical tests. The proteins associated with an adjusted p-value inferior to a FDR of 1% have been considered as significantly differentially abundant proteins. Finally, the proteins of interest are therefore those which emerge from this statistical analysis supplemented by those which are considered to be present from one condition and absent in another. Results of these differential analyses are summarized in Fig.3. For statistical analysis of output samples for a protein in order to be considered as a positive hit it needed to be detected in each of the three biological replicates. For total lysates (INPUTS) data analysis positive hits are proteins that were detected in at least two biological replicates.

### Constructing the Human Protein–Protein Interactome

We assembled 15 commonly used databases, focusing on high-quality PPIs with five types of evidences: (1) binary, physical PPIs tested by high-throughput yeast-two-hybrid (Y2H) screening system, combining binary PPIs tested from three publicly available high-quality Y2H datasets [56-58], (2) literature-curated PPIs identified by affinity purification followed by affinity-purification mass spectrometry (AP-MS), Y2H, and literature-derived low-throughput experiments; (3) binary, physical PPIs derived from protein three-dimensional structures; (4) kinase-substrate interactions by literature-derived low-throughput and high-throughput experiments; and (5) signaling networks by literature-derived low-throughput experiments. The protein-coding genes were mapped to their official gene symbols based on GeneCards (http://www.genecards.org/) and their Entrez ID. Computationally inferred interactions rooted in evolutionary analysis, gene expression data, and metabolic associations were excluded. The updated human interactome includes 243,603 PPIs connecting 16,677 unique proteins and is 40% greater in size compared to the previously used human interactome [59]. The human protein–protein interactome is provided in the Supplementary Data [60].

### PCA Based on the Split luciferase

HEK293T cells were seeded in white bottom 96-well flat microplate (#655083, Cellstar, Greiner Bio-One, Germany) at a concentration of 4.0×104 cells per well. After 24 h, cells were transfected using PEI-max (#24765-1, Polysciences) with 100 ng of pSNL-N2-C or pSNL-N2-Muc7 and 100 ng of pSNL-N1-”C interactors”. At 24 h post-transfection media was removed and Nluc enzymatic activity was measured using a Berthold Centro-XS luminometer by injecting 50 μL of luciferase substrate reagent (#E2820, Promega, France) per well and counting luminescence for 2s. Results were expressed as relative light units (RLU) or as a fold change normalized over the sum of controls, specified herein as normalized luciferase ratio (NLR). For a given protein pair A/B, NLR = (Nluc1-A + Nluc2-B)/[(Nluc1-A + Nluc2) + (Nluc1 + Nluc2-B)]. For the NLR validation experiment, each protein pair was assessed three times or more. The NLR of protein pair was considered as ‘validated’ if above a threshold value of 7.9 and above the upper limit of the confidence interval defined for the Muc7/bait pairs (Fig. 6*B*).

**Fig. 2.**
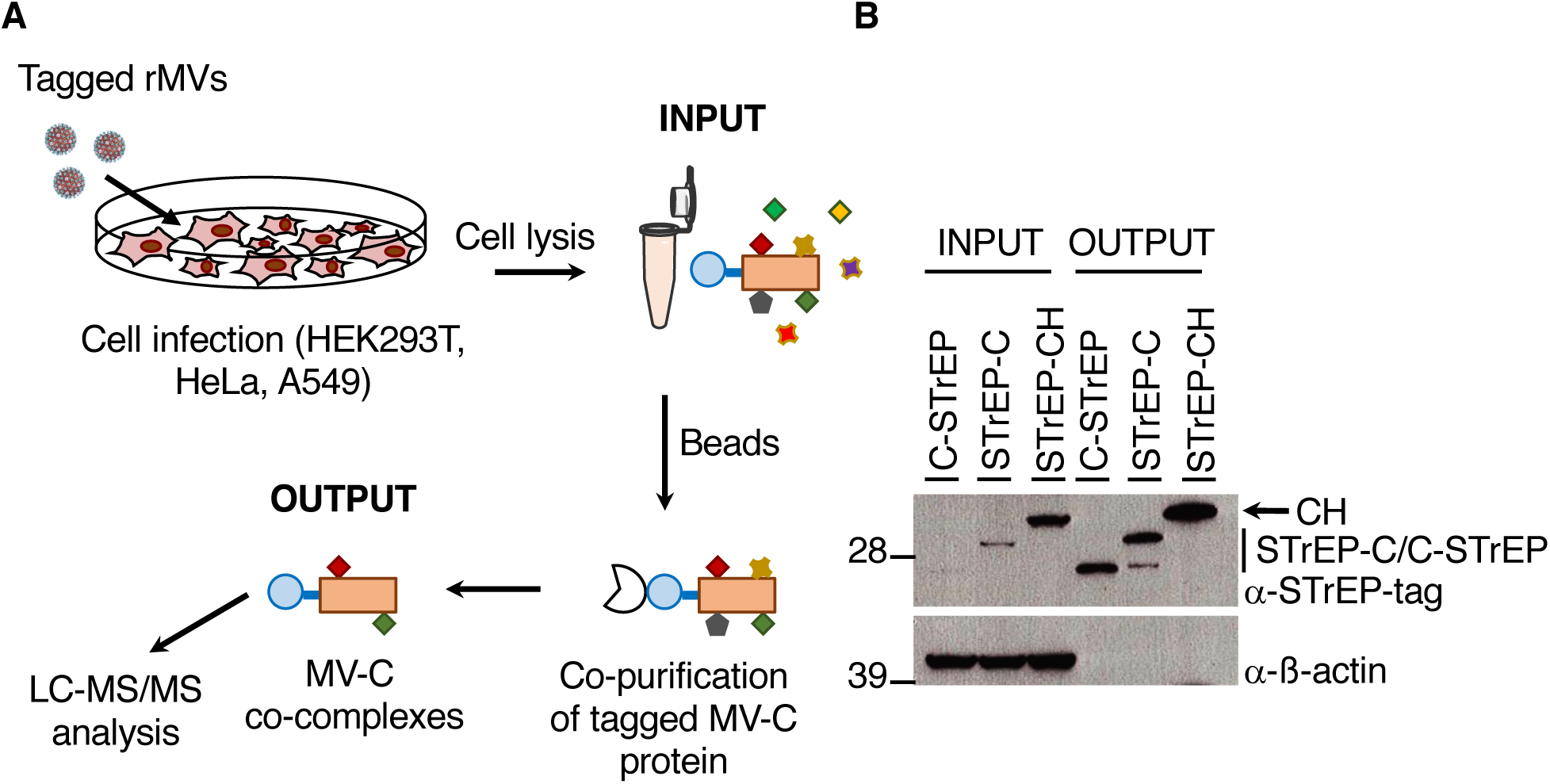
rMV3/STrEP-C and rMV3/C-STrEP recombinant viruses allow purification of MV-C-specific protein co-complexes. *A*. The protocol used to purify STrEP-C and associated cellular proteins from infected cell for MS-analysis of MV-C-specific protein factors. HEK293T, HeLa and A549 cells were infected with the corresponding recombinant tagged viruses. At 24 h post-infection, cells were lysed, and STrEP-C, C-STrEP, and STrEP-CH were co-purified with interacting cellular proteins using StrepTactin Sepharose beads. After two subsequent washing steps, protein complexes were released from the beads and prepared for LC-MS/MS. *B*. HEK293T cells were infected with either rMV3/STrEP-CH, or rMV3/STrEP-C, or rMV3/C-STrEP. Total lysates (INPUT) and MV-C specific protein complexes (OUTPUT) were analyzed by western blot using the One-STrEP-tag antibody. Western blot analysis of β-actin served as a control for loading and for nonspecific binding.

**Fig. 3.**
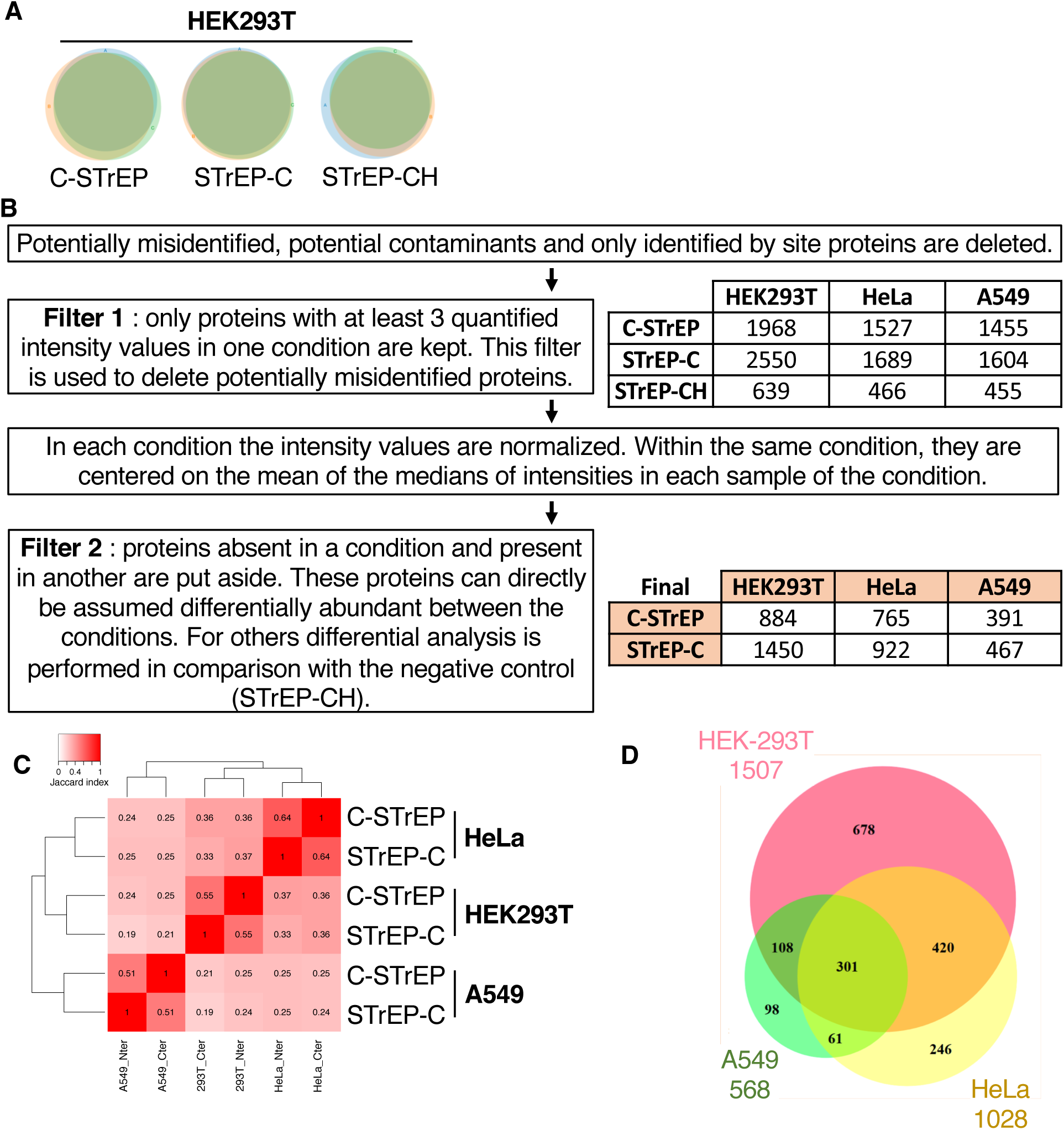
Mass spectrometry analysis of MV-C protein partners. *A*. Pie charts representing the protein overlap among the three different biological triplicates C-STrEP, STrEP-C, and STrEP-CH in HEK293T cells. *B*. Bioinformatics/statistical analysis pipeline from mass spectrometry data analysis. Tables show the number of identified proteins at each step of the analysis. *C*. Jaccard matrix showing the similarity of proteins isolated after purification with either C-STrEP or STrEP-C recombinant viruses in three different cell lines. The clustering is unsupervised and groups similar rows/columns together. *D*. Venn diagram representing final numbers and the overlap in MV-C-specific protein partners for HEK293T, HeLa, A549 cells.

**Fig. 4.**
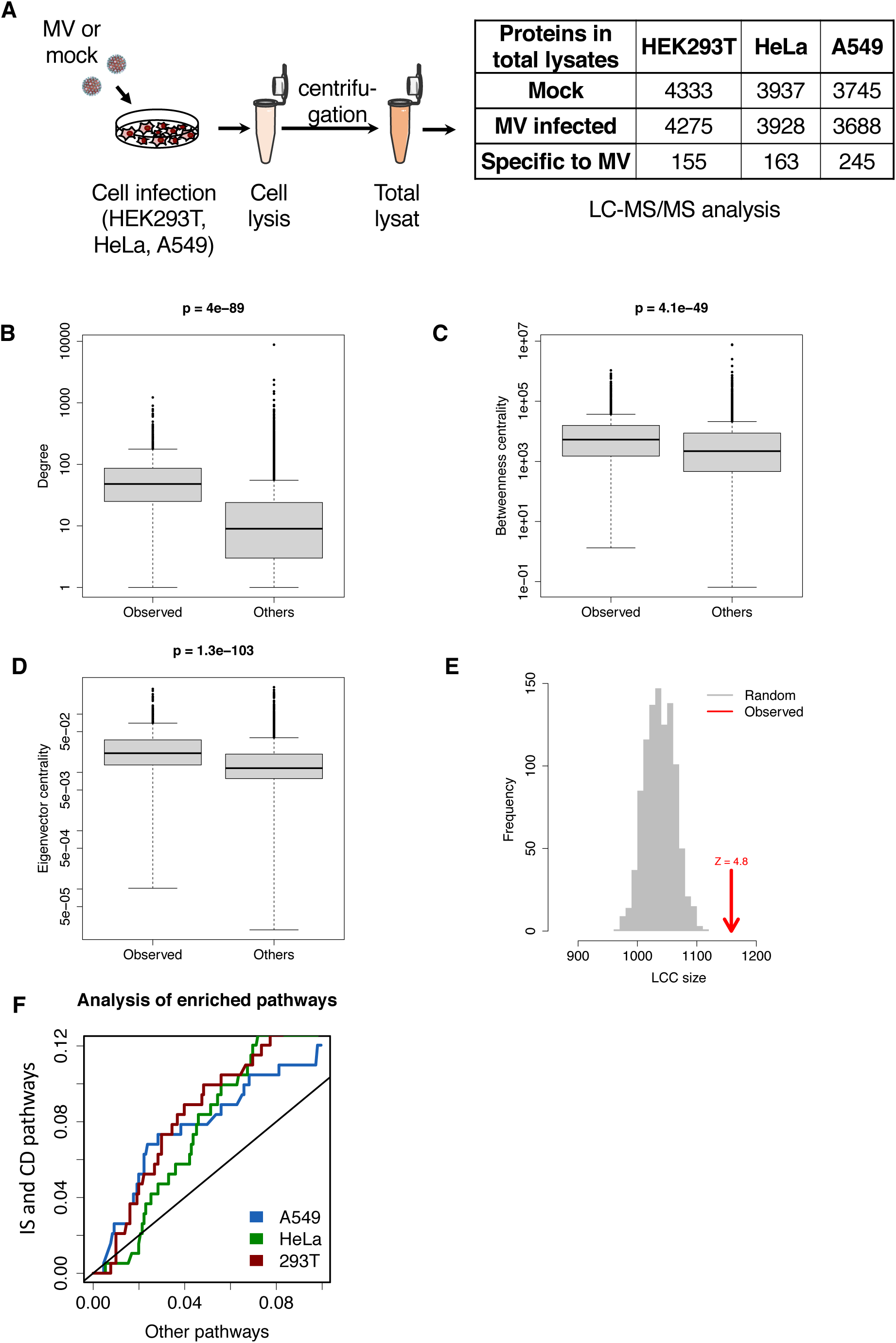
Topological analysis and cell pathways enrichment analysis of MV-C interaction network for HEK293T cell line. *A*. Representation of determination of total cell lysates. Cells were either mock infected or infected with Schwarz vaccine strain of MV. 24h later, same protocol of purification as in Figure 2 was applied. Number of proteins in total cell lysates *B*. Degree, *C*. betweenness and *D*. eigenvector centrality distributions of MV-C interaction partners and proteins from the rest of the interactome. *E*. Distribution of sizes of the largest connected component (LCC) of random sets of proteins with the same number as MV-C interaction partners in the interactome. The arrow shows the observed LCC for MV-C in HEK293T. *F*. ROC curves showing the enrichment in immune system (IS) and cell death (CD) pathways (overall pathways) in HEK293T, HeLa and A549 cells (see Supp.Table 4). Curves above the diagonal indicate enrichment in true positive pathways (IS and SD).

**Fig. 5:**
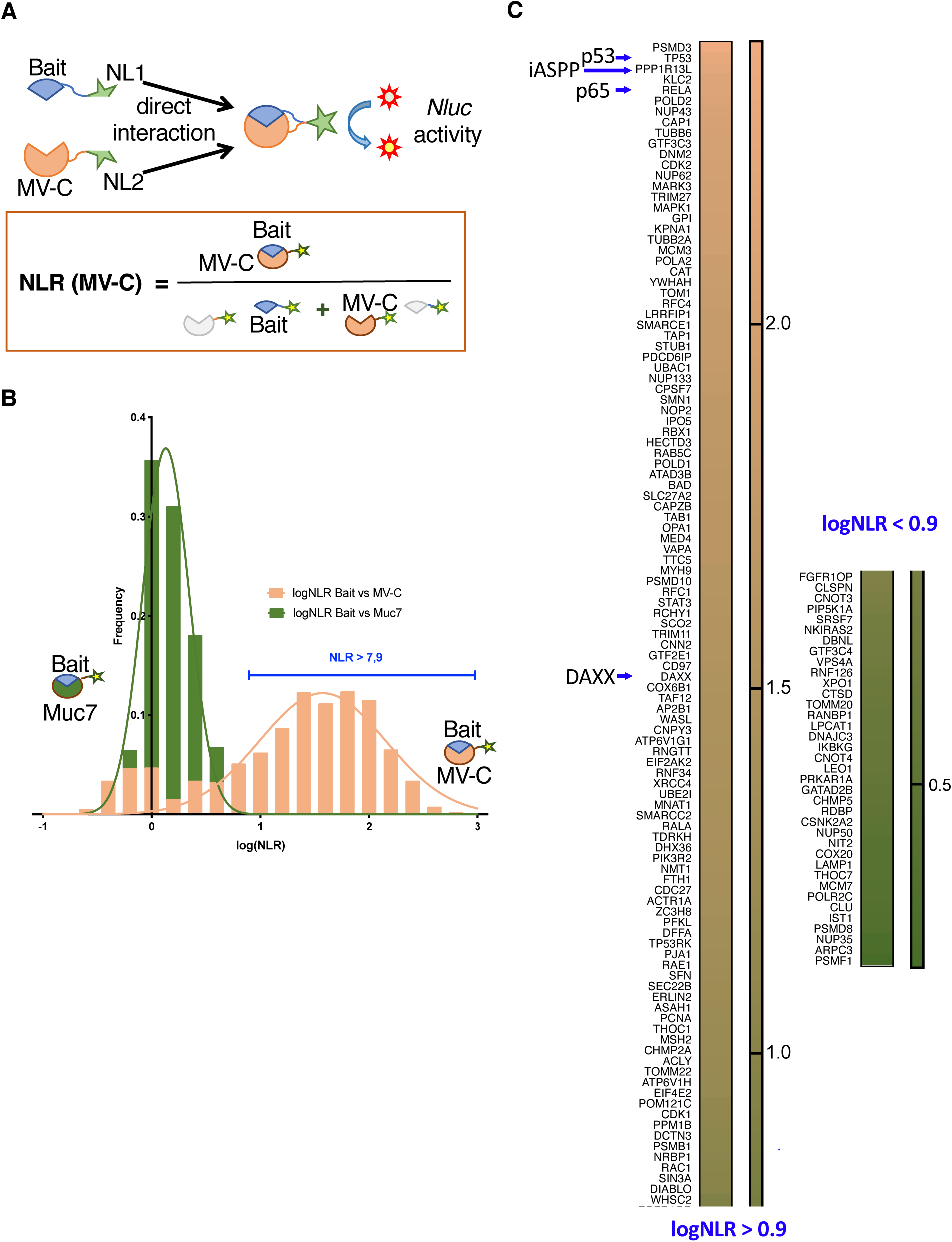
Protein complementation assay for MV-C protein and 146 host protein partners. *A*. Schematic representation of PCA approach. A host protein (prey) is fused downstream to the N-half of the Nluc (NL1). MV-C (bait) is fused downstream to the C-half of the Nluc (NL2). The formula to calculate Normalized Luciferase Ratio (NLR) for a given protein pair is shown. *B*. Frequency distribution of log(NLR) values for the MV-C and the Muc7 with corresponding superimposed fitted Gaussian curves. The positive threshold of 7.9 used in the network is indicated in green. Two biological replicates were performed for screening against Muc7 in order to establish the threshold. *C*. For the MV-C PCA, each protein pair was assessed in three biological replicates. The NLR of protein pair was considered as ‘validated’ if above the threshold value of 7.9.

**Fig. 6:**
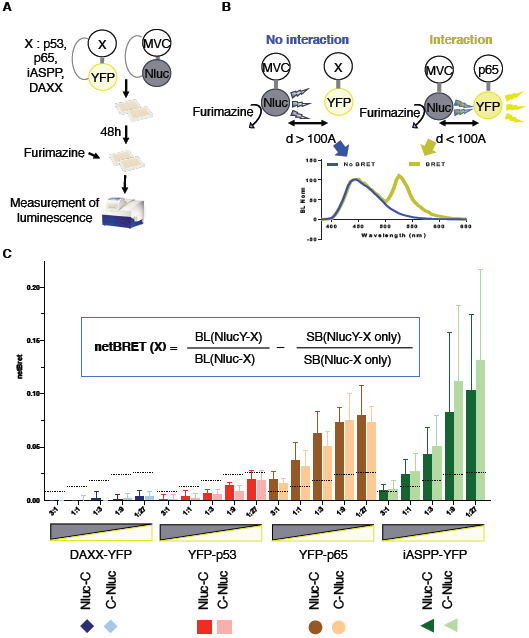
BRET analysis of MV-C direct interactions with DAXX, IASPP, p65 or p53. *A*. BRET method applied to study interactions between MV-C and DAXX, iASPP, p65 or p53. HEK293T cells were transfected for 48h with two plasmids expressing YFP- and Nluc-tagged proteins in N- and C-terminal positions at five DNA ratios, from 3:1 to 1:27. BRET was measured and 20 min later furimazine was added. Donor signal (Nluc, 448,5-472.5 nm emission) and acceptor signal (NlucY, 522.5-547.5nm emission) were measured using a bioluminescence plate reader. *B*. Molecular principle of the BRET assay. The donor, Nluc oxidizes furimazine substrate into Furimamide emitting light around 460nm. When BRET pair tagged proteins directly interact, proximity of Nluc will excite YFP, which in turn emits a yellow light around 530nm. If there is no interaction, no yellow emission will occur. *C*. Assessment of MV-C PPI with DAXX, iASPP, p65 or p53. Dot lines represent SOD1 versus either Nluc-C or C-Nluc level and is used as a negative control. Each value is the mean of all the combinations between Nluc-tagged host proteins and YFP-tagged viral proteins of each protein performed in three biological replicates with error bars representing the standard deviation. netBret represents the acceptor/donor ratio by dividing the initial fluorescence measurements (530 nm) by the luminescence measurements (485 nm) and subtracting out background luminescence (BRET readings in cells expressing the donor without any acceptor).

### Bioluminescence Resonance Energy Transfer

HEK293T were seeded in 384-well flat (4,000 cells/well) bottom µClear plates (#781097, Greiner Bio-One, Germany) coated with fibronectin (#354008, BD Biosciences, Germany). Transfection of HEK293T cells was performed at 80% confluence using 25ng of each plasmid construction mixed with 150nL of Fugene 6 (E2691, Promega, France) reagent per well. 24 h after transfection, fluorescence imaging measurements were performed using an Operetta widefield HCS system (Perkin Elmer, MA) equipped with a Peltier-cooled CCD camera with 1.3 megapixels per frame, 14-bit resolution, and a 20x long working distance objective lens with a 0.45 numerical aperture (NA). Then, media was removed and replaced by Dulbecco’s modified Eagle’s medium without phenol red (#21063029, Gibco, Switzerland), supplemented with the assay substrate Nano-Glo containing the furimazine at 200-fold dilution (#N1110, Promega, France).The bioluminescence signal of Nluc donor and the YFP acceptor emission were acquired using an EnVision Multimode Plate Reader (Perkin Elmer, USA) sequentially using a low and high band pass filter (460/25nm, 535/25nm), with an acquisition time of 0.5 sec. In addition, Emission spectral scans of the HEK293T cells expressing recombinant Nluc fusions positive and negative control were performed using a SpectraMax M5 fluorescence microplate reader (Molecular Devices, USA). Emission spectra were recorded from 350 nm to 700 nm using an integration time of 500 ms with 1nm step increments. All spectra were normalized to the luminescence value at the emission maximum (449 nm) of Nluc.

### Experimental Design and Statistical Rationale

For each experimental condition three biological replicates were performed and analysed each time. For mass spectrometry Cherry-infected cells were analysed as a control to remove all non-specific interactions (Fig.3*B*).

## RESULTS

### Generation and Characterization of Recombinant MV Expressing a Tagged MV-C

To co-purify MV-C with associated protein complexes from infected cells, we expressed MV-C from MV genome with a fusion tag allowing its capture. To reach this goal, we took advantage of reverse genetics for cloning and rescuing genetically modified MV [41] and of a high-throughput proteomic approach to study host protein partners of viral proteins in infected cells [10]. Both N- and C-terminal tags were considered. Sequences encoding the MV-C protein with N- or C-terminal One-STrEP tag were inserted into the pTM-MVSchw plasmid that contains a full-length infectious MV genome [41]. Expressing additional copies of MV-C from infectious MV appeared difficult. Indeed, expression from an additional transcription unit located downstream of the P gene (ATU-2) (Fig. 1*A*) was impossible after numerous attempts for either N- or C-terminally tagged C protein. Most rescued viruses either had stop codon that blocked the expression of the second copy of MV-C or introduced numerous mutations within MV-C coding sequence (data not shown). However, we succeeded in introducing an additional copy of MV-C gene with N- or C-terminal One-STrEP tag in ATU3 located downstream of the H gene (Fig. 1*A*). Recombinant viruses encoding One-STrEP-tagged MV-C protein were rescued and polyclonal population of each recombinant virus was obtained. Modified recombinant viruses were designated respectively rMV3/C-STrEP and rMV3/STrEP-C. As a negative control, we generated N-terminally tagged recombinant virus expressing the red fluorescent protein (CHERRY, CH) from ATU3 (designated rMV3/STrEP-CH).

To validate the expression of MV-C and CH-tagged proteins from rMV3/C-STrEP, rMV3/STrEP-C, and rMV3/STrEP-CH HeLa cells were infected with recombinant viruses at MOI of 1, and protein expression was determined 24 h post-infection by western blot analysis (Fig. 1B). Tagged MV-C and CH proteins were detected in infected cells using anti-Streptag antibody and anti-C antibody thus validating the expression of second copy of MV-C protein by rMV3/C-STrEP and rMV3/STrEP-C viruses (Fig. 1B). Transcription of additional STrEP-C and C-STrEP genes was also controlled by RT-PCR and Sanger sequencing analyses of total RNA extracted from rMV3/C-STrEP and rMV3/STrEP-C-infected cells. We observed that rMV3/C-STrEP was represented by a mixed recombinant virus population with approximately 60% corresponding to the rMV3/C-STrEP virus and other 40% recombinant MV possessed STOP codon at the beginning of C-STrEP sequence thus not interfering with STrEP-tag affinity purification. Our polyclonal rMV3/STrEP-C virus encoded STrEP-C with an additional adenine nucleotide resulting in the mutation of the last C-terminal 14 AA of C protein. Any additional attempts to obtain full length STrEP-C encoded by MV were unsuccessful. Due to the numerous problems to rescue absolutely perfect recombinant MV expressing additional copy of MV-C protein, we decided to perform our mass spectrometry-based MV-C interactomic analysis on two polyclonal populations of the rMV3/C-STrEP and rMV3/STrEP-C viruses described above, and to apply additional conventional approaches to further validate this high-throughput interactomic analysis.

We assessed the replication efficiency of One-STrEP-tagged MV recombinant viruses by determining single-step growth curves of rMV3/C-STrEP, rMV3/STrEP-C, rMV3/STrEP-CH and by comparing them to standard MV Schwarz (MVSchw). We infected Vero cells with the different One-STrEP-tagged viruses and MVSchw) at MOI 0.1. The growth of recombinant viruses was similar to that of unmodified MVSchw, and titers were comparable (Fig. 1*C*). To further test the potential impact of an additional copy of MV-C gene on viral replication, we compared the infection efficacy of HEK293T, HeLa, and A549 cells by the recombinant viruses at MOI 1 at 24 h post-infection. Immunostaining of MV nucleoprotein and flow cytometry analysis were used to detect infection (Fig. 1*D*). The efficient replication of rMV3/C-STrEP, rMV3/STrEP-C and rMV3/STrEP-CH viruses was comparable in the three cell lines. In conclusion, recombinant viruses expressing tagged MV-C protein or CH efficiently propagated with no detectable interference for viral replication.

### Identification of Host Protein Partners of MV-C by Affinity Purification and Shotgun Proteomic

To obtain a comprehensive list of direct and indirect interactors of MV-C protein, we applied a previously validated approach based on affinity purification coupled to mass spectrometry analysis of protein co-complexes [10]. To gain a larger spectrum of potential host protein partners of MV-C protein, the analysis was performed on three different cell lines: HEK293T, HeLa, and A549. Human Embryonic Kidney 293 cell line (HEK293) is extensively validated cell line for high-throughput methods [10, 35, 61-64]. A549 and HeLa cells are both human cancer cells: A549 (lung epithelial carcinoma) and HeLa (cervical epithelial adenocarcinoma) and are interesting to study considering the oncolytic activity of MV. The three cell lines were infected with rMV2/STrEP-C and rMV2/C-STrEP at MOI of 1. Tagged MV-C proteins were purified 24 h post-infection by affinity chromatography and co-purified cellular proteins (OUTPUT) were directly identified by nano-LC-MS/ MS analysis (Fig. 2*A*). The efficiency of purification was controlled by western blotting (Fig. 2*B*). The same experiment was repeated three times to obtain three biological replicates for each experimental condition. To distinguish between MV-C-associated proteins and non-specific binding to the beads, co-affinity purification experiments in HEK293T, HeLa, and A549 cells were performed in parallel with the control rMV3/STrEP-CH virus (Fig. 1*A*).

To analyze the results, we first filtered reverse proteins and potential contaminant proteins. We found large overlaps between the three biological replicates, demonstrating the robustness of our experimental approach (Fig. 3*A* and Supp.Fig. 1*A,B*, Supp.Table 2). Further MV-C-specific protein partners were determined by applying additional bioinformatic and statistical analyses (Fig.3*B*): only proteins with at least three quantified intensity values in one condition were kept. In each condition, the intensity values were normalized. Within the same condition, each protein intensity value was centered on the mean of the medians of intensities in each sample of the condition. Then, proteins absent in one condition but present in another were put aside and were directly assumed differentially abundant between the conditions. For other proteins, a differential analysis based on the LFQ values was performed for each MV-C protein interactor to compare with the negative control (STrEP-CH). Next, proteins that were found statistically relevant and exclusively present in MV-C sample were then assembled for each MV-C condition (Fig.3*B* final table). The similarity of proteins captured by MV-C N- and C-terminal tags was studied by calculating Jaccard index between EntrezIDs obtained in the three cell lines. We observed that each cell type naturally clustered together and resulted in 50-60% overlap for N-terminal and C-terminal MV-C (Fig. 3*C*). Due to this high number of host protein partners common for N- and C-terminally tagged MV-C the two sets of data for each cell line were assembled together and further compared. Although some cell-specific MV-C protein partners were found, more than 50% of captured proteins were shared between the three different cell lines (Fig.3*D*). A total of 1507, 1028 and 568 of specific protein partners for MV-C protein were specifically captured in HEK293T, HeLa, and A549 cell lines, respectively (Fig.3*D* and Supp.Table 2).

Within this list, some direct interactors of MV-C were previously identified by yeast two hybrid, including 26S proteasome regulatory subunit 6A (PSMC3) and WD repeat-containing protein 26 (WDR26) [29]. This confirms the sensitivity of our method. In contrast, we failed to identify SHCBP1 interaction with MV-C. Preferential localization of SHCBP1 to nucleus upon infection with the MVSchw in comparison to the minigenome [29] need to be assessed and might explain the failure to detect MV-C/SHCBP1 complex. Altogether, these data show that our mapping strategy is sensitive, and provides reproducible data.

### Topological and Pathway Analysis of MV-C /host Protein Interaction Network in HEK293T, HeLa and A549 Cells

We first established a specific list of total proteins for HEK293T, HeLa, and A549 cell lines. For this purpose, each cell line under study was either infected or not with MV Schwarz at MOI of 1 and lysed 24h post-infection. Total protein lysates (INPUTS) were extracted and subjected to shotgun proteomics. As previously mentioned, we performed bioinformatic and statistical analyses on total protein data and three cell-specific total protein lists were built (Fig.4*A*, Supp.Table *3*). We next analyzed MV-C protein interaction networks in HEK293T, A549, HeLa for specific topological features using several standard network centrality measures [65]. We first asked whether host proteins that co-purified with MV-C are central in the human interactome network [66]. Local (degree) and global (betweenness) centrality measures were calculated. The degree of a protein in a network corresponds to its number of direct partners and is, therefore, a measure of local importance. Betweenness is a global measure of centrality, as it measures the number of shortest paths (the smallest distances between any two proteins in the network) that go through a given protein. The degree distribution of cellular proteins co-purified with MV-C was plotted, and compared to the distribution obtained for the rest of the human interactome network (Fig. 4*B*, Supp.Figs.2.*A*, 3.*A*). These distributions were significantly distinct (p-value < 4e x 10-89 under a Mann Whitney test for HEK293T for HEK293T). Thus, the cellular proteins that co-purified with MV-C exhibit much more cellular partners than normally expected by chance. We also calculated and plotted the betweenness centrality distribution for the cellular proteins that co-purified with MV-C (Fig. 4*C*, Supp.Figs.2.*B*, 3.*B*). This distribution was significantly different from the one obtained for the randomized human interactome network (p-value < 4.1x 10-49 for HEK293T). Thus, betweenness centrality measures show that MV-C interacting partners are enriched for proteins that connect multiple regions in the human interactome. Furthermore, we measured the eigenvector centrality of MV-C host protein partners. This approach measures the influence of a node in a network. Relative scores are assigned to all nodes in the network based on the concept that connections to high-scoring nodes contribute more to the score of the node in question than equal connections to low-scoring nodes. For MV-C protein network we observed nodes were connected to many nodes who themselves had high scores (Fig. 4*D*, Supp.Figs.2.*C*, 3.*C*). Next, we explored whether MV-C interacting proteins were forming cohesive subgraphs, as is generally observed for “disease modules” [59]. For this, we computed the size of the largest connected component (LCC) of the subgraph formed by the MV-C host protein partners. We then selected the same number of proteins at random in the interactome and computed the size of the largest connected component, repeating the process 1,000 times. Finally, we computed a Z-score to quantify the significance of the observed value, corresponding to the number of standard deviations that the observed value departed from the expected value of the random distribution. We found for HEK293T a Z-score of 4.8 (N=1,158 proteins out of 1,498 in the LCC, compare to 1,045±23 expected at random), showing that MV-C proteins form a cohesive disease module (Fig. 4*E*, Supp.Figs.2.*D*, 3.*D*).

Finally, we tested the hypothesis that the two major cellular pathways which are **I**mmune **S**ystem and **C**ell **D**eath pathways (overall referred to as IS and CD) were specifically enriched in the MV-C network for the three cell types (Supp.Table 5, 6 and 7). For this, we performed a pathways enrichment analysis on MV-C specific networks in the three different cell lines. Similarly to the topological analysis, the total protein dataset was used as a background for all pathways and disease gene enrichments (Fig.4*A*, Supp.Table *3*). Pathways were taken from MSigDB v3.1 [67] and Wikipathways Kelder, 2012 #1979}, with a total of 8690 sets of genes. For each pathway, the significance of its overlap with MV-C partners was assessed by computing a Hypergeometric p-value, with a background set to be the total protein dataset. To test that IS and CD pathways were overrepresented in the most enriched pathways, we computed a Receiver Operating Characteristic (ROC) curve for each cell type (Fig. 4*F*). We ranked pathways according to their p-value, from lowest to highest. We then plotted the ROC curves, showing the enrichment in IS and CD pathways (y axis, true positive rate) in the top of the list compare to other pathways (x axis, false positive rate) and showed the enrichment for the three cell types (Fig. 4*F*). Altogether, topological analyses of MV-C interaction network demonstrate that MV-C preferentially interacts, either directly or indirectly, with host proteins that are central in the human interactome network. MV-C protein co-complex represents well defined functional modules as assessed by the enrichment for proteins sharing interactions. This is clearly in the line of proposed role of MV-C in regulation of innate immunity response and cells death pathways.

### Protein Complementation Assay (PCA) to Study MV-C Direct Protein Partners Involved in Immune Response and Cell Death Pathways

Based on the results of our pathway enrichment analysis, we selected 146 MV-C-specific protein partners (Supp.Table 1). To further specify PPIs between MV-C and the 146 selected host protein partners involved in cell death, immune response and infectious disease pathways we applied a protein complementation assay PCA based on the split luciferase [68]. We used a modified version of the PCA approach assigned NlucPCA for Nanoluc luciferase PCA (N2H, [43]). In NlucPCA, the luciferase activity is restored leading to luminescence signal when the two fragments of the Nluc are in close contact (10 nm) resulting from the direct PPI between the prey and the bait fused to Nluc luciferase fragments 1 and 2 (NL1 and NL2), respectively (Fig.5.*A*). 146 potential interactors (Supp.Table 1) were screened against MV-C. These 146 host proteins were screened against Muc7 protein used as a negative control to determine the cut-off of non-specific binding. The Gaussian repartition of the normalized luminescence ratio (NLR) of Muc7 and MV-C gave us a threshold at an NLR of 7.9 (Fig.5. *B*). From the list of 146 MV-C specific protein partners, we validated the direct interaction with MV-C for 109 host proteins (Fig.5. *C*). Thus, PCA provided additional validation to sensitivity and specificity of our mass-spectrometry approach and further narrowed the specific list of MV-C protein network to its direct PPIs.

### Bioluminescence Resonance Energy Transfer (BRET) Method Confirms MV-C Direct PPIs with iASPP and p65

We found high NlucPCA scores for p53, iASPP, and p65 (Fig.5.*C*) and further addressed their direct interactions with MV-C. Additionally, we included DAXX protein to this list as it has earlier been reported to be targeted by RNA viruses to control proapoptotic genes expression [69]. None of the selected proteins (DAXX, p65, p53, and IASPP) have previously been described as direct partner of MV-C. We first infected HeLa or HEK293T cells with recombinant viruses expressing tagged MV-C to validate by affinity chromatography purification coupled to western blot analysis the interaction of MV-C with p65, IASPP, and p53 in an infection context (Supp.Fig.4). However, we failed to validate MV-C binding to DAXX (data not shown).

To study MV-C PPIs directly in live cells with simultaneous control of protein expression efficacy, we applied the BRET approach. To this aim, DAXX, iASPP, p53, and p65 were fused with YFP coding sequence, and MV-C was tagged with Nluc either N-terminal or C-terminal (Fig. 6.*A*). BRET signals indicating MV-C interaction with either DAXX, iASPP, p53, or p65 were measured in living cells 48h post-transfection of plasmids expressing YFP- and Nluc-tagged proteins at different concentration ratios of DNA. As positive and negative controls, we used EYFP fused to Nluc and superoxide dismutase 1 (SOD1) fused with the Nluc C-terminally, respectively. For each protein pair, we calculated the NetBret which represents the direct bioluminescence from the donor (Nluc) and the acceptor (NlucY) normalized to the SOD1-Nluc control (Fig. 6.*B,C*). We observed that for every protein under study the netBret value increased while the expression of donor increase (Fig. 6.*C*) and thus the ratio MV-C:interactor decreased. The NetBret values were similar for MV-C tagged C- or N-terminally with the Nluc. Thus, similar to tagged protein affinity purification results (Fig. 3*C*), the interaction was not influenced by the tag and both N- and C-terminal tagged MV-C interacted at the same level. MV-C protein interactions with iASPP and p65 presented significantly high value of netBret, whereas MV-C interaction with p53 and DAXX demonstrated low values. Thus, we failed to validate p53 and DAXX direct PPI with MV-C (Fig. 6.*C*).

## DISCUSSION

This study aimed to establish the MV-C protein network of interactions in MV-infected cells. We used a strategy based on recombinant viruses expressing tagged viral proteins followed by affinity purification and a bottom-up mass spectrometry-based proteomics [10]. This high-throughput approach allowed us to obtain a list of MV-C specific host partners interacting *via* direct or indirect PPIs. By performing topological and signaling pathway-oriented analyses of MV-C host protein partners we observed that MV-C was embedded in a complex host interactome. Advanced computational tools recently developed in network biology highlighted the immunity and cell death pathways as specifically enriched in the MV-C network for the three cell types tested: HEK293T, HeLa, A549 (Fig.4.*F*). Further, we mainly focused on these two cell pathways, as MV vaccine and modified recombinant MV strains, are of high application in vaccinology and in cancer treatment.

In order to validate and to specify PPIs found by high-throughput MS approach, we applied two unbiased approaches to study PPIs: PCA based on the split luciferase and BRET. PCA allowed us to validate that from a list of 146 host proteins that form a specific protein co-complex with MV-C upon infection and belong to cell death, immune response and infectious disease pathways, 109 proteins were direct interactors (Fig.5.*C*). Thus, the PCA provided additional validation to sensitivity and specificity of our mass-spectrometry approach and further narrowed the MV-C protein network to its direct PPIs.

p53, iASPP, and p65 were three host factors that drew our attention as they possessed the highest luciferase scores in PCA (Fig.5.*C*). Together with DAXX, their interactions with MV-C were further studied by BRET approach. None of these proteins (DAXX, p65, p53, and iASPP) has previously been described as a direct partner of MV-C. p65, p53 and iASPP proteins interact with each other. Indeed, iASPP inhibits p53-mediated apoptosis [70, 71] and binds RelA/p65 to inhibit its transcriptional activity [72, 73]. These two binding partners of iASPP explain why under certain conditions iASPP may have pro- and anti-apoptotic effects, depending on iASPP regulation of p53 or RelA/p65, respectively. High netBret values confirmed MV-C protein interactions with iASPP and p65 (Fig.6.*C*). However, we failed to validate MV-C interaction with p53 by BRET analysis. Indeed, p53 demonstrated low netBret values that were comparable with the negative control (SOD1). As no single PPI assay is exceptionally superior to any other, the results of PCA and BRET analyses should be combined to maximize PPIs detection. In addition, we validated MV-C specific PPI with iASPP, p53 and RelA/p65 in infected cells (Supp.Fig.4). Our study demonstrates that iASPP, p53 and RelA/p65 proteins are in complex with MV-C in MV-infected cells. Further studies should question whether MV-C binding with these three proteins promotes an anti-apoptotic or a pro-apoptotic role of iASPP. Previous studies also suggested that MV-C binding to p65 could suppress NF-kB promoter activity [74]. However, additional analyses are needed in the future to understand the functional role of MV-C interaction with iASPP, p53, and RelA/p65.

We also included DAXX in our BRET analysis of interactions with MV-C, although its luciferase score was lower than for iASPP, p53, and RelA/p65. DAXX has previously been reported to be targeted by the mutant M protein of the oncolytic Vesicular Stomatitis Virus (VSV, order *Mononegavirales*), and this PPI is implied in the induction of apoptosis [69]. However, in our study BRET approach and affinity chromatography purification of MV-C protein co-complexes from infected cells failed to validate MV-C specific binding to DAXX. Thus, additional PPI assays should be applied to examine MV-C PPI with DAXX.

Viral virulence factors are embedded into a complex host interactome, enabling the virus to control the host cell to provide efficient viral replication. MV-C is such an example. We applied various PPI approaches that allowed us to glimpse the interplay of the dual functional role of MV-C in controlling both cell death and host innate immune responses. We believe that our results provide a first solid support for further studies that will focus on precise molecular mechanisms by which MV-C controls of the host cells.

## ABBREVIATIONS

AA: Amino Acid
ATU: Additional Transcription Unit
BRET: Bioluminescence Resonance Energy Transfer
FDR: False Discovery Rate
h: hour
IKKα: IkB Kinase α
IFN: interferon
JAK: JAnus Kinase
RIG-I: Retinoic acid-Inducible Gene-I
LCC: Largest Connected Component
LGP2: Laboratory of Genetics and Physiology 2
MAVS: Mitochondrial AntiViral-Signaling protein
MDA5: Melanoma Differentiation-Associated protein 5
min: minute
MOI: Multiplicity Of Infection
MV: Measles Virus
NES: Nuclear Export Signal
NLR: Normalized Luminescence Ration
NLS: Nuclear Localization Signal
Nluc: Nanoluciferase
NF-kB: Nuclear Factor-kappa B
PCA: Protein Complementation Assay
PKR: Protein Kinase R
PPI: Protein-Protein Interaction
RIG-I: Retinoic acid-Inducible Gene-I
RLU: Relative Luciferase Unit
s: second
STAT1: Signal Transducer and Activator of Transcription 1
STAT2: Signal Transducer and Activator of Transcription 2
TRIM25: TRIpartite Motif containing protein 25
*wt*: wild type

## Acknowledgments

We thank Dr. Kaoru Takeuchi and Dr. Chantal Rabourdin-Combe for anti-C rAb polyclonal antibody and anti-N mAb monoclonal antibody, respectively. We thank Dr.Hervé Bourhy from Lyssavirus Epidemiology and Neuropathology Laboratory of Institut Pasteur for providing us pYFP-p65 plasmid which contains EYFP reporter gene fused to p65. We thank all members of the Viral Genomics and Vaccination Unit and Laboratory of Molecular Genetics of RNA Viruses for their critical discussion and Dr. Atousa Arbabian for technical support.

## DATA AVAILABILITY

The mass spectrometry proteomics data have been deposited to the ProteomeXchange Consortium *via* the PRIDE [75] partner repository with the dataset identifier PXD015316. * This work was supported by ANR 16 CE18 0016 01 Tangy ONCOMEVAX, the Institut Pasteur and CNRS. AM was supported by IDF DIM 2016.

## REFERENCES

1. Frantz, P.N., et al., Measles-derived vaccines to prevent emerging viral diseases. Microbes Infect, 2018. 20(9-10): p. 493–500.

2. Russell, S.J., et al., Oncolytic Measles Virotherapy and Opposition to Measles Vaccination. Mayo Clin Proc, 2019.

3. Cattaneo, R., et al., Measles virus editing provides an additional cysteine-rich protein. Cell, 1989. 56(5): p. 759–764.

4. Bellini, W.J., et al., Measles virus P gene codes for two proteins. J Virol, 1985. 53(3): p. 908–919.

5. Caignard, G., et al., Measles virus V protein blocks Jak1-mediated phosphorylation of STAT1 to escape IFN-alpha/beta signaling. Virology, 2007. 368(2): p. 351–362.

6. Caignard, G., et al., Differential regulation of type I interferon and epidermal growth factor pathways by a human Respirovirus virulence factor. PLoS Pathog, 2009. 5(9): p. e1000587.

7. Jiang, Y., et al., Host-Pathogen Interactions in Measles Virus Replication and Anti-Viral Immunity. Viruses, 2016. 8(11).

8. Mandhana, R., et al., Constitutively Active MDA5 Proteins Are Inhibited by Paramyxovirus V Proteins. J Interferon Cytokine Res, 2018. 38(8): p. 319–332.

9. Cruz, C.D., et al., Measles virus V protein inhibits p53 family member p73. J Virol, 2006. 80(11): p. 5644–5650.

10. Komarova, A.V., et al., Proteomic analysis of virus-host interactions in an infectious context using recombinant viruses. Mol Cell Proteomics, 2011. 10(12): p. M110 007443.

11. Sanchez-Aparicio, M.T., et al., Paramyxovirus V Proteins Interact with the RIG-I/TRIM25 Regulatory Complex and Inhibit RIG-I Signaling. J Virol, 2018. 92(6).

12. Nagai, Y., et al., Accessory genes of the paramyxoviridae, a large family of nonsegmented negative-strand RNA viruses, as a focus of active investigation by reverse genetics. Curr Top Microbiol Immunol, 2004. 283: p. 197–248.

13. Alkhatib, G., et al., Expression of bicistronic measles virus P/C mRNA by using hybrid adenoviruses: levels of C protein synthesized in vivo are unaffected by the presence or absence of the upstream P initiator codon. J Virol, 1988. 62(11): p. 4059–4069.

14. Nishie, T., et al., The C protein of wild-type measles virus has the ability to shuttle between the nucleus and the cytoplasm. Microbes Infect, 2007. 9(3): p. 344–354.

15. Devaux, P., et al., Attenuation of V-or C-defective measles viruses: infection control by the inflammatory and interferon responses of rhesus monkeys. J Virol, 2008. 82(11): p. 5359–5367.

16. Escoffier, C., et al., Nonstructural C protein is required for efficient measles virus replication in human peripheral blood cells. J Virol, 1999. 73(2): p. 1695–1698.

17. McAllister, C.S., et al., The RNA-activated protein kinase enhances the induction of interferon-beta and apoptosis mediated by cytoplasmic RNA sensors. J Biol Chem, 2009. 284(3): p. 1644–1651.

18. Mrkic, B., et al., Lymphatic dissemination and comparative pathology of recombinant measles viruses in genetically modified mice. J Virol, 2000. 74(3): p. 1364–1372.

19. Nakatsu, Y., et al., Translational inhibition and increased interferon induction in cells infected with C protein-deficient measles virus. J Virol, 2006. 80(23): p. 11861–11867.

20. Patterson, J.B., et al., V and C proteins of measles virus function as virulence factors in vivo. Virology, 2000. 267(1): p. 80–89.

21. Pfaller, C.K., et al., Measles Virus Defective Interfering RNAs Are Generated Frequently and Early in the Absence of C Protein and Can Be Destabilized by Adenosine Deaminase Acting on RNA-1-Like Hypermutations. J Virol, 2015. 89(15): p. 7735–7747.

22. Pfaller, C.K., et al., Measles virus C protein impairs production of defective copyback double-stranded viral RNA and activation of protein kinase R. J Virol, 2014. 88(1): p. 456–468.

23. Richetta, C., et al., Sustained autophagy contributes to measles virus infectivity. PLoS Pathog, 2013. 9(9): p. e1003599.

24. Takeuchi, K., et al., Stringent requirement for the C protein of wild-type measles virus for growth both in vitro and in macaques. J Virol, 2005. 79(12): p. 7838–7844.

25. Fontana, J.M., et al., Regulation of interferon signaling by the C and V proteins from attenuated and wild-type strains of measles virus. Virology, 2008. 374(1): p. 71–81.

26. Shaffer, J.A., et al., The C protein of measles virus inhibits the type I interferon response. Virology, 2003. 315(2): p. 389–397.

27. Yokota, S., et al., Measles virus C protein suppresses gamma-activated factor formation and virus-induced cell growth arrest. Virology, 2011. 414(1): p. 74–82.

28. Sparrer, K.M., et al., Measles virus C protein interferes with Beta interferon transcription in the nucleus. J Virol, 2012. 86(2): p. 796–805.

29. Ito, M., et al., Measles virus nonstructural C protein modulates viral RNA polymerase activity by interacting with host protein SHCBP1. J Virol, 2013. 87(17): p. 9633–9642.

30. Nakatsu, Y., et al., Measles virus circumvents the host interferon response by different actions of the C and V proteins. J Virol, 2008. 82(17): p. 8296–8306.

31. Pfaller, C.K., et al., The C protein is recruited to measles virus ribonucleocapsids by the phosphoprotein. J Virol, 2019.

32. Mura, M., et al., Non-encapsidated 5’ copy-back defective-interfering genomes produced by recombinant measles viruses are recognized by RIG-I and LGP2 but not MDA5. J Virol, 2017.

33. McAllister, C.S., et al., Mechanisms of protein kinase PKR-mediated amplification of beta interferon induction by C protein-deficient measles virus. J Virol, 2010. 84(1): p. 380–386.

34. Runge, S., et al., In vivo ligands of MDA5 and RIG-I in measles virus-infected cells. PLoS Pathog, 2014. 10(4): p. e1004081.

35. Sanchez David, R.Y., et al., Comparative analysis of viral RNA signatures on different RIG-I-like receptors. Elife, 2016. 5.

36. Hur, S., Double-Stranded RNA Sensors and Modulators in Innate Immunity. Annu Rev Immunol, 2019. 37: p. 349–375.

37. Xia, M., et al., Mitophagy enhances oncolytic measles virus replication by mitigating DDX58/RIG-I-like receptor signaling. J Virol, 2014. 88(9): p. 5152–5164.

38. Toth, A.M., et al., Protein kinase PKR mediates the apoptosis induction and growth restriction phenotypes of C protein-deficient measles virus. J Virol, 2009. 83(2): p. 961–968.

39. Itoh, M., et al., Increased induction of apoptosis by a Sendai virus mutant is associated with attenuation of mouse pathogenicity. J Virol, 1998. 72(4): p. 2927–2934.

40. Koyama, A.H., et al., Virus multiplication and induction of apoptosis by Sendai virus: role of the C proteins. Microbes Infect, 2003. 5(5): p. 373–378.

41. Combredet, C., et al., A molecularly cloned Schwarz strain of measles virus vaccine induces strong immune responses in macaques and transgenic mice. J Virol, 2003. 77(21): p. 11546–11554.

42. Calain, P., et al., The rule of six, a basic feature for efficient replication of Sendai virus defective interfering RNA. J Virol, 1993. 67(8): p. 4822–4830.

43. Choi, S.G., et al., Maximizing binary interactome mapping with a minimal number of assays. Nat Commun, 2019. 10(1): p. 3907.

44. Besson, B., et al., Regulation of NF-kappaB by the p105-ABIN2-TPL2 complex and RelAp43 during rabies virus infection. PLoS Pathog, 2017. 13(10): p. e1006697.

45. Kim, J., et al., Nanoluciferase signal brightness using furimazine substrates opens bioluminescence resonance energy transfer to widefield microscopy. Cytometry A, 2016. 89(8): p. 742–746.

46. Takeuchi, K., et al., Measles virus V protein blocks interferon (IFN)-alpha/beta but not IFN-gamma signaling by inhibiting STAT1 and STAT2 phosphorylation. FEBS Lett, 2003. 545(2-3): p. 177–182.

47. Giraudon, P., et al., Monoclonal antibodies against measles virus. J Gen Virol, 1981. 54(Pt 2): p. 325–332.

48. Erde, J., et al., Enhanced FASP (eFASP) to increase proteome coverage and sample recovery for quantitative proteomic experiments. J Proteome Res, 2014. 13(4): p. 1885–1895.

49. Cox, J., et al., MaxQuant enables high peptide identification rates, individualized p.p.b.-range mass accuracies and proteome-wide protein quantification. Nat Biotechnol, 2008. 26(12): p. 1367–1372.

50. Cox, J., et al., Andromeda: a peptide search engine integrated into the MaxQuant environment. J Proteome Res, 2011. 10(4): p. 1794–1805.

51. Cox, J., et al., Accurate proteome-wide label-free quantification by delayed normalization and maximal peptide ratio extraction, termed MaxLFQ. Mol Cell Proteomics, 2014. 13(9): p. 2513–2526.

52. Wieczorek, S., et al., DAPAR & ProStaR: software to perform statistical analyses in quantitative discovery proteomics. Bioinformatics, 2017. 33(1): p. 135–136.

53. Smyth, D.J., et al., A genome-wide association study of nonsynonymous SNPs identifies a type 1 diabetes locus in the interferon-induced helicase (IFIH1) region. Nat Genet, 2006. 38(6): p. 617–619.

54. Ritchie, M.E., et al., limma powers differential expression analyses for RNA-sequencing and microarray studies. Nucleic Acids Res, 2015. 43(7): p. e47.

55. Giai Gianetto, Q., et al., Calibration plot for proteomics: A graphical tool to visually check the assumptions underlying FDR control in quantitative experiments. Proteomics, 2016. 16(1): p. 29–32.

56. Luck, K., et al., A reference map of the human binary protein interactome. Nature, 2020. 580(7803): p. 402–408.

57. Rolland, T., et al., A proteome-scale map of the human interactome network. Cell, 2014. 159(5): p. 1212–1226.

58. Rual, J.F., et al., Towards a proteome-scale map of the human protein-protein interaction network. Nature, 2005. 437(7062): p. 1173–1178.

59. Menche, J., et al., Disease networks. Uncovering disease-disease relationships through the incomplete interactome. Science, 2015. 347(6224): p. 1257601.

60. Sun, X., et al., High-throughput methods for combinatorial drug discovery. Sci Transl Med, 2013. 5(205): p. 205rv201.

61. Baum, A., et al., Differential recognition of viral RNA by RIG-I. Virulence, 2011. 2(2): p. 166–169.

62. Chiang, J.J., et al., Viral unmasking of cellular 5S rRNA pseudogene transcripts induces RIG-I-mediated immunity. Nat Immunol, 2018. 19(1): p. 53–62.

63. van der Veen, A.G., et al., The RIG-I-like receptor LGP2 inhibits Dicer-dependent processing of long double-stranded RNA and blocks RNA interference in mammalian cells. EMBO J, 2018. 37(4).

64. Zhao, Y., et al., RIG-I like receptor sensing of host RNAs facilitates the cell-intrinsic immune response to KSHV infection. Nat Commun, 2018. 9(1): p. 4841.

65. Jalili, M., et al., Evolution of Centrality Measurements for the Detection of Essential Proteins in Biological Networks. Front Physiol, 2016. 7: p. 375.

66. Cheng, F., et al., Network-based prediction of drug combinations. Nat Commun, 2019. 10(1): p. 1197.

67. Liberzon, A., et al., Molecular signatures database (MSigDB) 3.0. Bioinformatics, 2011. 27(12): p. 1739–1740.

68. Dixon, A.S., et al., NanoLuc Complementation Reporter Optimized for Accurate Measurement of Protein Interactions in Cells. ACS Chem Biol, 2016. 11(2): p. 400–408.

69. Gaddy, D.F., et al., Oncolytic vesicular stomatitis virus induces apoptosis via signaling through PKR, Fas, and Daxx. J Virol, 2007. 81(6): p. 2792–2804.

70. Bergamaschi, D., et al., iASPP oncoprotein is a key inhibitor of p53 conserved from worm to human. Nat Genet, 2003. 33(2): p. 162–167.

71. Samuels-Lev, Y., et al., ASPP proteins specifically stimulate the apoptotic function of p53. Mol Cell, 2001. 8(4): p. 781–794.

72. Trigiante, G., et al., ASPP [corrected] and cancer. Nat Rev Cancer, 2006. 6(3): p. 217–226.

73. Yang, J.P., et al., NF-kappaB subunit p65 binds to 53BP2 and inhibits cell death induced by 53BP2. Oncogene, 1999. 18(37): p. 5177–5186.

74. Schuhmann, K.M., et al., The measles virus V protein binds to p65 (RelA) to suppress NF-kappaB activity. J Virol, 2011. 85(7): p. 3162–3171.

75. Perez-Riverol, Y., et al., The PRIDE database and related tools and resources in 2019: improving support for quantification data. Nucleic Acids Res, 2019. 47(D1): p. D442-D450.

